# Canopy position has a stronger effect than tree species identity on phyllosphere bacterial diversity in a floodplain hardwood forest

**DOI:** 10.1101/2020.02.07.939058

**Authors:** Martina Herrmann, Patricia Geesink, Ronny Richter, Kirsten Küsel

**Affiliations:** Institute of Biodiversity, Aquatic Geomicrobiology, Friedrich Schiller University Jena, Dornburger Strasse 159, D-07743 Jena, Germany; German Center for Integrative Biodiversity Research, Deutscher Platz 5e, 04103 Leipzig, Germany; Systematic Botany and Functional Biodiversity, Institute for Biology, Leipzig University, Johannisallee 21, 04103 Leipzig; Geoinformatics and Remote Sensing, Institute of Geography, Johannisallee 19a, Leipzig University, 04103 Leipzig, Germany

**Keywords:** phyllosphere, canopy crane, *Acer pseudoplatanus*, *Quercus robur*, *Tilia cordata*

## Abstract

The phyllosphere is a challenging microbial habitat in which microorganisms can flourish on organic carbon released by plant leaves but are also exposed to harsh environmental conditions. Here, we assessed the relative importance of canopy position – top, mid, and bottom at a height between 31 m and 20 m – and tree species identity for shaping the phyllosphere microbiome in a floodplain hardwood forest. Leaf material was sampled from three tree species - maple (*Acer pseudoplatanus L.*), oak (*Quercus robur L.*), and lime (*Tilia cordata MILL.*) - at the Leipzig canopy crane facility (Germany). Estimated bacterial species richness (Chao1) and bacterial abundances approximated by quantitative PCR of 16S rRNA genes exhibited clear vertical trends with a strong increase from the top to the mid and bottom position of the canopy. 30 Operational Taxonomic Units (OTUs) formed the core microbiome, which accounted for 77% of all sequence reads. These core OTUs showed contrasting trends in their vertical distribution within the canopy, pointing to different ecological preferences and tolerance to presumably more extreme conditions at the top position of the canopy. Co-occurrence analysis revealed distinct tree species-specific OTU networks, and 55-57% of the OTUs were unique to each tree species. Overall, the phyllosphere microbiome harbored surprisingly high fractions of Actinobacteria of up to 46%. Our results clearly demonstrate strong effects of the position in the canopy on phyllosphere bacterial communities in a floodplain hardwood forest and - in contrast to other temperate or tropical forests - a strong predominance of Actinobacteria.

## Introduction

The phyllosphere is an important microbial habitat which spans about 10^8^ km^2^ on a global scale (Lindow and Brandl 2003). It is host to central biogeochemical processes as well as plant-microbe-interactions that affect plant community dynamics, and ecosystem functioning and productivity (Vorholt 2012, Laforest-Lapointe & Whitaker 2019). Microbiota on leaf surfaces contribute to biogeochemical processes such as N_2_-fixation (Freiberg 1998, Moyes et al. 2016), nitrification (Guerrieri et al. 2015), and transformation of C1 compounds or terpenes and monoterpenes released from the plants (Bringel and Coue 2015). Especially in forest canopies, they may play a central role in the bioremediation of air pollutants (Wei et al. 2017) and are in exchange with atmospheric and cloud microbiota, suggesting important implications for climate regulation (Bringel and Coue 2015). Beyond their role in biogeochemical processes, phyllosphere microbiota are not only passive inhabitants on surfaces of plants but interact with their host in multiple ways (Dees et al. 2015), resulting in plant-microbe relationships that range from loose associations to defined symbioses (Bringel and Coue 2015). They produce phytohormones or affect the production of these hormones by the plant (Brandl & Lindow 1998, Gourion et al. 2006, Reed et al. 2010) and they improve host resistance against pathogens (Innerebner et al. 2011, Balint-Kurti et al. 2010).

However, the phyllosphere is also a harsh environment where microorganisms are exposed to extreme conditions such as high UV radiation, desiccation, rainfall, antimicrobial substances released by leaves, and strong nutrient limitation (Bringel and Coué 2015). Altogether, these environmental parameters positively select for the bacterial taxa that are able to persist on leaves (Knief et al. 2010). As a consequence, phyllosphere bacterial diversity has been shown to be much lower than diversity of the rhizosphere, soil or marine ecosystems (Delmotte et al., 2009; Knief et al., 2012, Haas et al. 2018). Especially in the canopy of large trees, selective environmental forces that restrict microbial diversity are likely to vary along vertical gradients (Laforest-Lapointe et al. 2016a) with the severest stress by abiotic parameters presumably acting at the top of the canopy. However, studies addressing intra-individual variability of phyllosphere microbial communities have so far mostly focused on a single tree species such as *Gingko biloba* or *Magnolia grandiflora* (Leff et al. 2015, Stone and Jackson 2019) or on rather small trees at a maximum height of 6 m (Laforest-Lapointe et al. 2016a). Ongoing colonization of leaves by microorganisms and their continuous removal, e. g., by rain fall, result in complex community dynamics in the forest canopy phyllosphere (Bringel and Coue 2015). However, it is unclear if microbial taxa differ in their preference for a particular canopy position and how this translates into the spatial heterogeneity of canopy-associated biogeochemical processes. Despite their high relevance for ecosystem functioning, phyllosphere microbiota in forest canopies have so far received comparatively little attention. Factors such as host species identity, leaf age, location in the canopy, light incidence, and microclimate conditions have been identified as central factors shaping the phyllosphere environment in tropical and North American temperate forests (Lambais et al., 2006; Yadav et al., 2005, Redford et al. 2010, Kembel et al. 2014, Laforest-Lapointe et al. 2016b, 2017, 2019). However, it is unclear if the phyllosphere diversity patterns observed for tropical and North American forests, especially the strong effect of host species identity, also apply to European forests with a different tree species composition

Here, we hypothesize that (i) forest trees harbor species-specific phyllosphere bacterial communities, and that (ii) microbial communities at the treetop are the most distinct, as they are the most exposed to abiotic stress factors. Taking advantage of the Leipzig canopy crane facility located in central Germany, allowing us to sample leave material from up to 33 m height, we compared phyllosphere microbial communities between three different tree species abundant in the Leipzig floodplain forest – *A. pseudoplatanus L., Q. robur* L., and *T. cordata* MILL. - and across three different height levels within the canopy - top, mid, and bottom. Our results revealed clear vertical trends of increasing bacterial diversity and abundances, and changes in community structure from the top of the canopy to mid and bottom canopy, which were further modulated by plant species identity.

## Methods

### Leipzig floodplain hardwood forest and canopy crane facility

Leaf samples were obtained from three tree species – *Q. robur L.* (oak; Qr), *A. pseudoplatanus L.* (maple; Ap), and *T. cordata MILL.* (lime; Tc) - in the Leipzig floodplain hardwood forest, located near the city of Leipzig in Germany (Fig. 1a). Situated in the floodplain of the Elster, Pleiße and Luppe rivers, the Leipzig floodplain forest is one of the largest floodplain forests in Central Europe (Müller 1995). Climatic conditions are characterized by warm summers and an annual mean temperature of 8.4°C with an annual precipitation of 516 mm (Jansen 1999). The forest consists of the ash-elm floodplain forest (*Fraxino-Ulmetum*) and is dominated by maple (*A. pseudoplatanus L.)*, ash (*Fraxinus excelsior L.)*, oak (*Q. robur L.)*, and hornbeam *(Carpinus betulus L.)*, with smaller contribution of lime (*T. cordata MILL.)* and elm *(Ulmus minor MILL.)* (Otto and Floren 2010). A crane facility (Leipzig Crane facility, LCC) for the investigation of forest tree canopies was established in this floodplain forest in 2001, allowing access to about 800 tree individuals in up to 33 m height, covering a total area of 1.65 ha (Fig. 1A). Different positions within the tree canopy were accessed by using a gondola attached to the crane. We sampled leaf material from the top, mid, and bottom position of the tree canopy. Depending of the height of individual trees, these positions ranged from 27.0-30.7 m, 18.6-26.3 m, and 12.9-23.2 m, respectively (Fig. 1b).

**Fig. 1.**
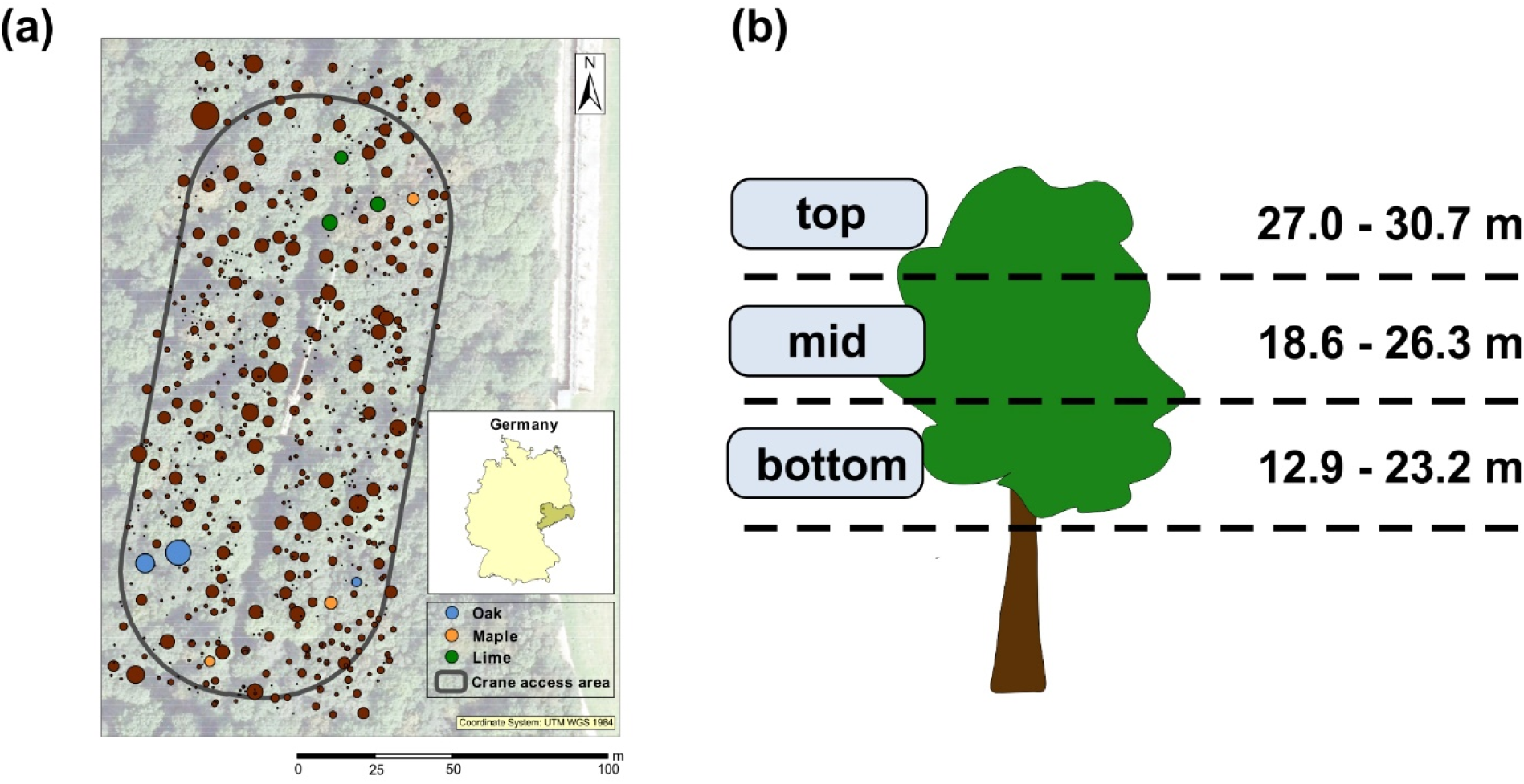
Study site and sampling design. (a) Location of the tree individuals sampled in this study within the total canopy crane research site. Tree species are distinguished by color. (b) Sampling design. Samples were taken in triplicates from the top, mid and bottom position of the canopy.

### Sampling of leaf material and detachment of surface-associated microbes

Leaves were sampled by clipping off leaves with ethanol-cleaned scissors, followed by immediate transfer to autoclaved polyphenylene ether (PPE) containers. Between five and ten leaves were sampled per tree individual, canopy position, and spatial replicate. Leaves were stored at ambient temperature (approx. 15°C) during transport and were immediately processed upon arrival at the laboratory within two hours. Leaves were amended with 250 ml suspension buffer (0.15 M NaCl, 1 % Tween 20) in the autoclaved containers in which leaves had been sampled, subjected to mild sonication (1 min at 10% intensity, turned and another 1 min at 10% intensity), followed by shaking for 20 min at 100 rpm at room temperature. Subsequently, suspensions were filtered through 0.2 µm polyethersulfone filters (Supor, Pall Corporation), and filters were stored at −80°C until nucleic acid extractions were performed. The remaining leaf material was dried at 50°C for one week for determination of dry weight.

### Nucleic acid extraction, Illumina MiSeq amplicon sequencing, and quantitative PCR

DNA was extracted from the filters using the DNEasy PowerSoil Extraction kit (Qiagen) following the manufacturer’s protocol. Filters were cut into smaller pieces to facilitate cell disruption during the bead-beating step. Amplicon sequencing of bacterial 16S rRNA genes was carried out targeting the V3-V4 region with the primer combination Bakt_0341F/Bakt_0785R (Klindworth et al. 2013). PCR amplification, library preparation, and sequencing on an Illumina MiSeq platform using v3 chemistry was performed at LGC (Berlin) as previously described (Rughöft et al. 2016). Abundances of bacterial 16S rRNA genes were determined by quantitative PCR using Brilliant SYBR Green II Mastermix (Agilent Technologies) on a Mx3000P system (Agilent Technologies) and the primer combination Bac8Fmod (Nercessian et al. 2005) and Bac338Rabc (Loy et al. 2002) as previously described (Herrmann et al. 2012). Due to loss of plant material of some samples before determination of leaf dry weight, abundance data are only available for a subset of all samples (see Supplementary Table 1).

### Sequence analysis

Sequence analysis was carried out using Mothur (Schloss et al. 2009) following the Schloss MiSeq SOP (Kozich et al. 2013) as previously described (Rughöft et al. 2016) along with the SILVA taxonomy reference database v132 (Quast et al. 2013). Chimera search was performed using the uchime algorithm implemented in Mothur. Operational Taxonomic Units (OTUs) were assigned on a 0.03 distance level using the vsearch algorithm. We obtained 6,037,303 high quality sequence reads across 81 samples with read numbers per sample ranging from 1610 to 198790. For further statistical analysis, read numbers were normalized to the same number for all samples (11188 reads) using the sub.sample function implemented in Mothur, resulting in the exclusion of two samples with too low read numbers from the data set (Ap6 and Qr18). Sequences obtained in this study have been submitted to the European Nucleotide Archive (ENA) under the study accession number PRJEB36420, sample accession numbers SAMEA6502636 – SAMEA6502715.

### Statistical analysis

Principal Component Analysis (PCA), PERMANOVA analysis, generation of Box and Whisker Plots, linear regression analysis, and pairwise Mann-Whitney U-test were carried out using the software PAST (Hammer et al. 2001). For PCA analysis, only OTUs with at least 10 reads across all samples were included. Co-occurrence network analysis was done using the MENA platform (Deng et al. 2012). Networks were constructed for the phyllosphere microbiome of each tree species, including samples from all three canopy positions, tree individuals and spatial replicates per tree species. Only OTUs with more than 20 sequence reads across all samples per tree species were included. Network calculations were based on Spearman rank correlation coefficients without log transformation of the data. Networks were graphically refined using Cytoscape 3.7.2.

## Results

### Effect of canopy position and tree species on phyllosphere bacterial communities

Amplicon sequencing of bacterial 16S rRNA genes revealed clear vertical trends in OTU richness and community composition from the top to the bottom canopy position, further modulated by the tree species. The number of observed and estimated (Chao estimator) species-level OTUs tended to increase from the top of the canopy towards the mid position (Fig. 2a). For the phyllosphere of lime, the increase in bacterial OTU numbers was already visible between the mid and the bottom position of the canopy while such a trend was less obvious for the other two tree species. Median values of observed (estimated) OTU numbers ranged from 264 to 331 (607 to 655) at the top of the canopy, from 332 to 400 (729 to 937) at the mid position, and from 393 to 429 (875 to 933) at the bottom of the canopy with the highest numbers observed in association with maple.

**Fig. 2.**
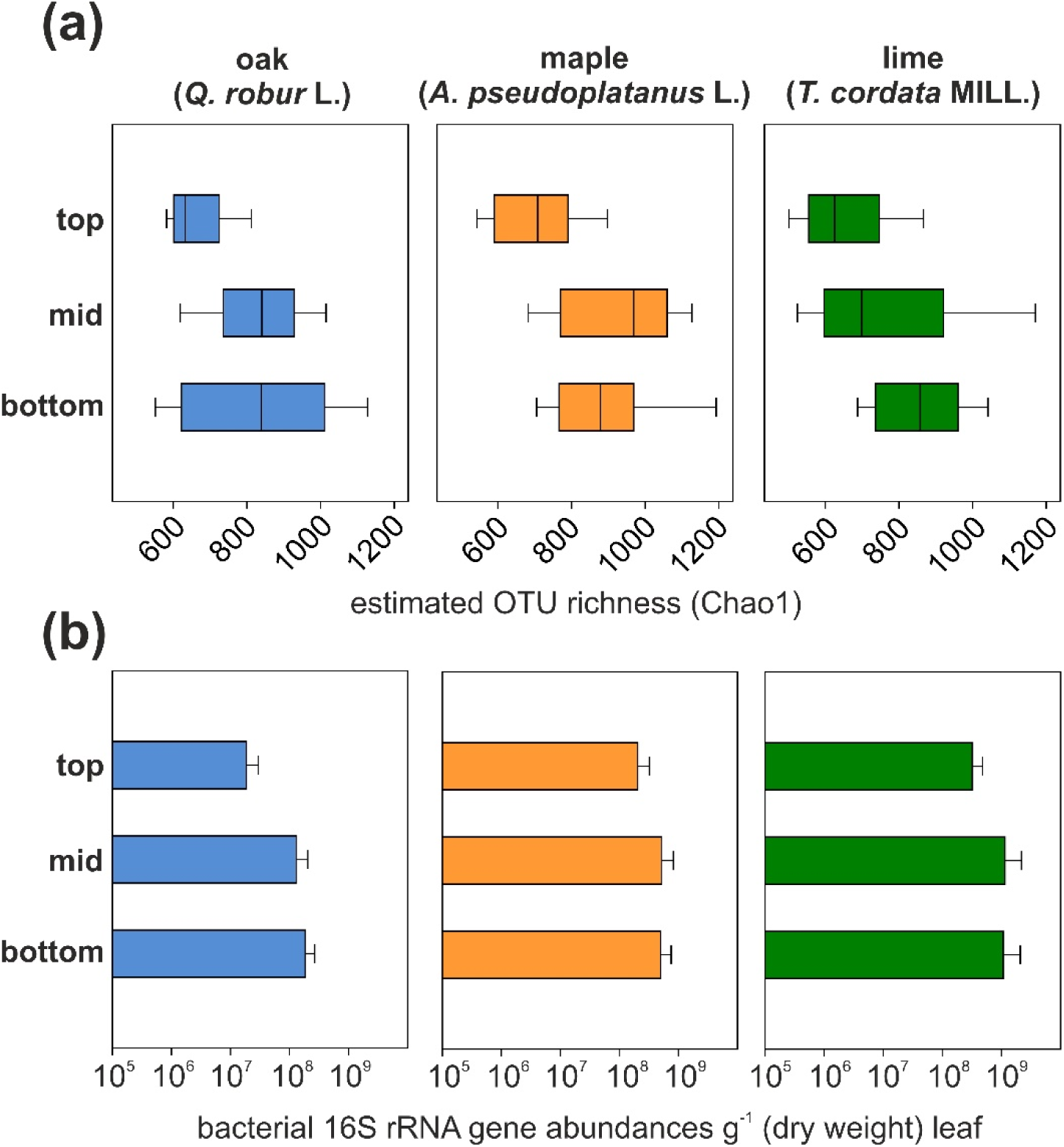
(a) Estimated bacterial species richness and (b) abundance of bacterial 16S rRNA genes per g leaf (dry weight) in the canopy of *Q. robur L.* (left panel), *A. pseudoplatanus L.* (mid panel), and *T. cordata MILL.* (right panel) with positions in the canopy categorized as “top”, “mid”, and “bottom”. Data are means (± standard deviation) of results obtained from three tree individuals with three replicates per sampled canopy area. For gene abundances, a reduced number of samples is available (see Supplementary Table 1).

Abundances of bacterial 16S rRNA genes per g dry weight leaf showed a similar trend as bacterial OTU richness, with lower abundances at the top of the canopy compared to the mid or bottom position. However, these differences were only significant for the oak phyllosphere and rather represent trends, as for some of the samples, especially from the top of the canopy, only a reduced number of replicates is available. Across all tree species, individuals, and position in the canopy, bacterial 16S rRNA gene abundances ranged from 6.8 × 10^6^ to 3.2 × 10^9^ g^−1^ (dry weight) (Fig. 2b). The lowest abundances were observed in association with oak and the highest in association with lime.

In line with these findings, PERMANOVA analysis based on Euclidean distance revealed that position in the canopy as well as tree species had a significant effect on phyllosphere bacterial community structure (p = 0.0001) with canopy position explaining 15% and tree species identity explaining 11.6% of the total community variation (Fig. 3b). In addition, interactions between these two factors also had a significant effect (p = 0.0059). The observed height- and tree species-dependent trends in bacterial OTU richness were reflected by clustering patterns of samples using Principal Component Analysis. Here, especially for lime and maple, samples obtained from the top of the canopy clustered separate from samples obtained from mid or bottom positions with clear differences in OTU composition between these two tree species (Fig. 3a). In contrast, for the mid and bottom position of the canopy, we observed minor clustering of communities according to tree species but also large overlaps. For each tree species, communities of the mid and bottom position were more similar to each other than they were to the communities at the top of the canopy. In fact, for all three tree species together, the fraction of OTUs shared between the top and mid canopy position (15 – 19.4%) or the top and bottom canopy position (15.5 – 20.5%) was significantly lower than the fraction of OTUs shared between the mid and bottom position (19.4 - 25.1%; Mann-Whitney-U-test, p = 0.00172) (Supplementary Fig. 1).

**Fig. 3.**
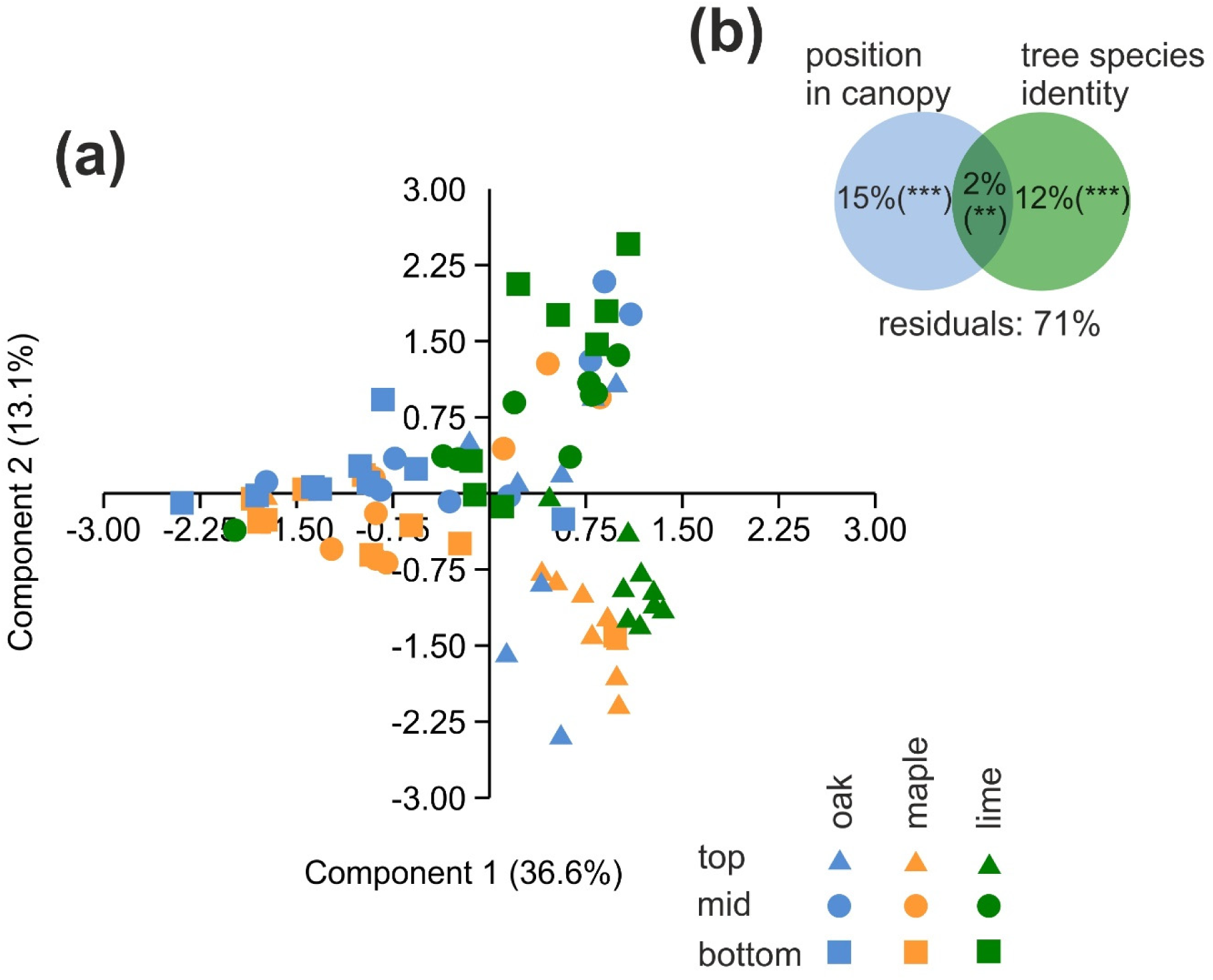
(a) Principal component analysis of phyllosphere bacterial communities across three tree species and top, mid, and bottom position of the canopy. (b) Variation partitioning resulting from PERMANOVA analysis. Analyses were based on distribution patterns of species-level OTUs using Euclidean distances. Colors denote tree species, symbols denote position within the tree canopy.

### Composition of the hardwood forest canopy microbiome

The phyllosphere bacterial communities associated with all three tree species were largely dominated by Actinobacteria, Bacteroidetes, Alphaproteobacteria, and Gammaproteobacteria which together accounted for at least 95% of the sequence reads in each sample (Fig. 4). Members of Deinococcus-Thermus, candidate phylum FBP, representatives of the Candidate Phyla Radiation (Hug et al 2016) such as *Cand*. Saccharimonadia, and Deltaproteobacteria were consistently present in the phyllosphere communities but rarely reached relative abundances of more than 3%. Taxa associated with chemolithoautotrophic lifestyles within the Gammaproteobacteria, such as *Nitrosomonas, Nitrosospira*, or *Ferribacterium*, were only represented by a few sequence reads across all samples.

**Fig. 4.**
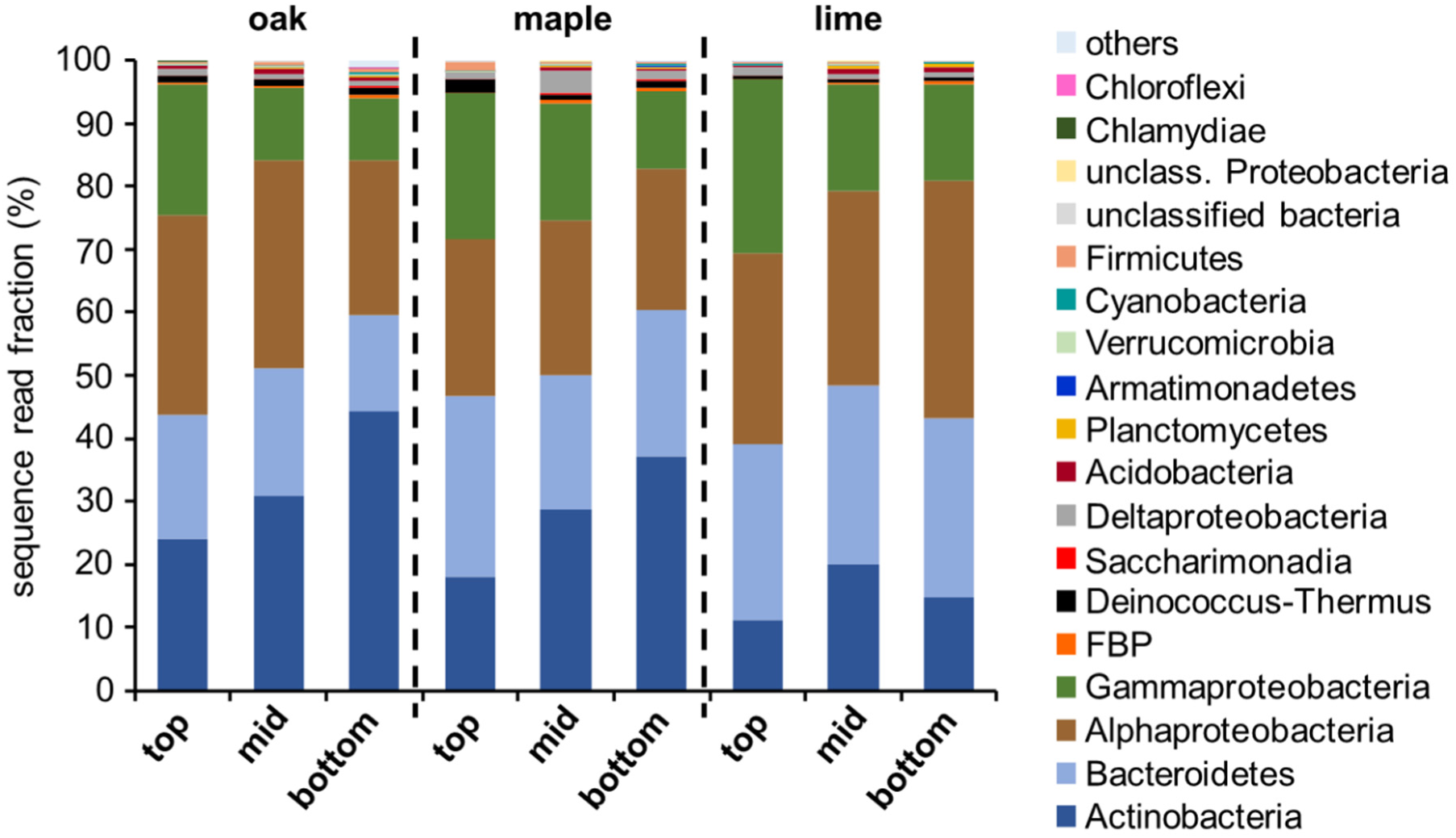
Composition of the bacterial communities associated with the phyllosphere of oak (*Q. robur L.)*, maple (*A. pseudoplatanus L.)*, and lime (*T. cordata MILL.)* at the top, mid and bottom position of the canopy. Each bar represents mean values of three tree individuals and three spatial replicates per tree individual. Taxonomic affiliation is shown on the phylum level or class level for *Proteobacteria* and *Patescibacteria*.

Notably, we observed major shifts in the relative fractions among the four dominant phyla in dependence of canopy position. For both maple and oak, the fraction of Actinobacteria increased strongly from the top of the canopy (18 and 24%, respectively) to 37 and 44% at the bottom of the canopy (p <0.006) while no such height-dependent trend was visible in association with lime trees (p = 0.55375) (Figure 4; Supplementary Fig. 2). Moreover, relative abundances of Actinobacteria were lower in the lime phyllosphere compared to maple and oak for the bottom position of the canopy (pairwise Mann-Whitney-U test; p = 0.008 and p = 0.006, respectively) and, compared to oak, also for the top position of the canopy (p = 0.004). In turn, the relative fraction of Gammaproteobacteria mostly represented by *Burkholderiacaea, Enterobacteriacaea, Diplorickettsiacaea*, and *Pseudomonadaceae* tended to decrease from the top towards the bottom of the canopy for all three tree species (Figure 4), however, these trends were not significant (Supplementary Fig. 2). Bacteroidetes, represented mostly by the families of *Hymenobacteraceae* and *Spirosomacea*e, did not show any obvious changes in relative abundance with canopy position on the phylum level.

Following changes in relative abundances of the most abundant 20 OTUs across the top, mid, and bottom position of the canopy, further demonstrated that the distribution patterns of individual OTUs were often linked to canopy position, which was further modulated by tree species identity. The strongest increase in relative abundance from the top towards the canopy mid and bottom was observed for OTU01 affiliated with *Friedmaniella* (*Propionibacteriaceae*), which constituted a dominant member of the phyllosphere community, accounting for up to 46% of the sequence reads in the individual samples at the canopy bottom and mid position but on average only for 4 – 11% at the top of the canopy (Supplementary Fig. 3). In turn, several OTUs decreased in relative abundance towards the mid and bottom position of the canopy for all three tree species, e. g., OTU09 (*Massilia*), OTU10 (*Hymenobacter*), OTU15 (*Methylobacterium*), and OTU17 (*Kineococcus*).

In the next step, we subjected relative abundances of these 20 OTUs across all samples to hierarchical clustering to further identify their preferential association with canopy position or a particular tree species. In general, clustering patterns according to tree species appeared to be less pronounced than those according to canopy position and confirmed the preferential association of OTU01 with bottom and mid canopy positions of oak and maple and its clear distinction from the distribution patterns of the other abundant OTUs (Fig. 5). OTU02, OTU03, OTU04, OTU05, OTU06, OTU07, OTU09 affiliated with *Beijerinckiaceae, Sphingomonadaceae, Hymenobacter*, and *Massilia* showed distribution patterns complementary to those of OTU01. OTU05 (*Sphingomonas*), OTU07 (*Hymenobacter*) and OTU09 (*Massilia*) occurred primarily in association with the canopy top position across all three tree species.

**Fig. 5.**
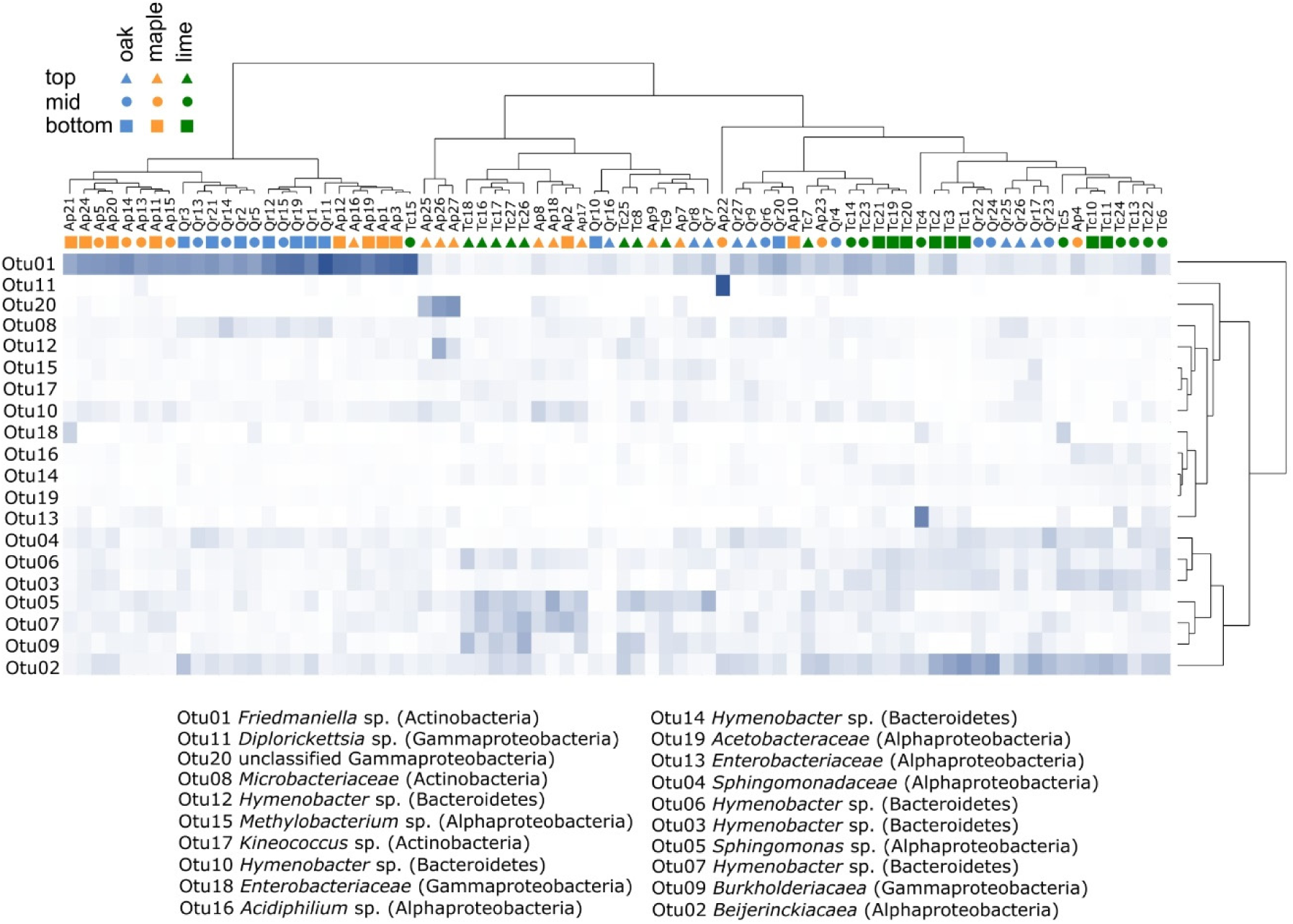
Relative abundance of the 20 most abundant species-level OTUs across all samples. Affiliation with top, mid or bottom position of the canopy is depicted by triangles, circles or squares, respectively. Names of samples refer to microbial communities in association with *A. pseudoplatanus L.* (Ap), *Q. robur L.* (Qr), and *T. cordata MILL.* (Tc). Two-way hierarchical clustering was performed using Euclidean distance.

### Phyllosphere core microbiome and co-occurrence networks

To further investigate the effect of tree species on the phyllosphere bacterial communities, we merged all the OTUs observed in association with a given tree species in the different samples to one OTU pool and compared the resulting three tree species-dependent OTU pools to each other. For each tree species, about 55-57% of the OTUs were unique to that tree species, while a fraction of 26-29% was shared between all three tree species. Maple and oak shared a slightly higher fraction of OTUs between their phyllosphere microbiomes (36-37%) compared to the fraction shared with lime (33 - 34%) (Supplementary Fig. 4).

30 species-level OTUs from four different phyla and 13 different families were present across all tree species and individuals, canopy positions, and spatial replicates, forming the phyllosphere core microbiome. Altogether, these 30 OTUs accounted for 77% of the sequence reads but only for 0.3% of the observed phyllosphere bacterial diversity, indicating that the phyllosphere communities were strongly dominated by these core microbiome representatives. Among the core microbiome members, *Hymenobacteraceae* (*Cytophagales, Bacteroidetes*) contributed the largest number of OTUs, followed by *Burkholderiaceae* (*Betaproteobacteriales*, Gammaproteobacteria) and *Beijerinckiacaea* (*Rhizobiales*, Alphaproteobacteria).

In the next step, we analyzed the interconnection of these core microbiome OTUs within the microbial communities associated with each tree species. Communities were subjected to co-occurrence network analysis, which revealed substantially different networks for each tree species (Fig. 6). We obtained networks with 156 nodes and 332 links for maple, 216 nodes and 258 links for oak, and 161 nodes and 365 links for lime. Notably, co-occurrence networks of the oak phyllosphere microbiomes showed the largest fraction of negative interactions (44.2%), while negative interactions accounted for only 11.1 or 20.3%, respectively, of all interactions in the phyllosphere OTU network of maple and lime. Across samples, OTU01 was not only the OTU with the highest relative abundance but also among the top five OTUs with the highest number of links to other OTUs within the maple and oak canopy. In contrast, OTU01 was less strongly connected in the phyllosphere of lime, coinciding with its lower relative abundance in association with that tree species. Overall, OTU01 exhibited mostly positive links to other OTUs. However, these associated OTUs differed substantially across tree species. For the oak phyllosphere, 63% of the OTUs associated with OTU01 were also *Actinobacteria*, while Proteobacteria dominated the associated OTUs in the lime phyllosphere. In association with maple, OTU01 exhibited the most diverse connections to other OTUs, including an especially high contribution of *Spirosomacea* and *Saccharimonadales* compared to the other two tree species.

**Fig. 6.**
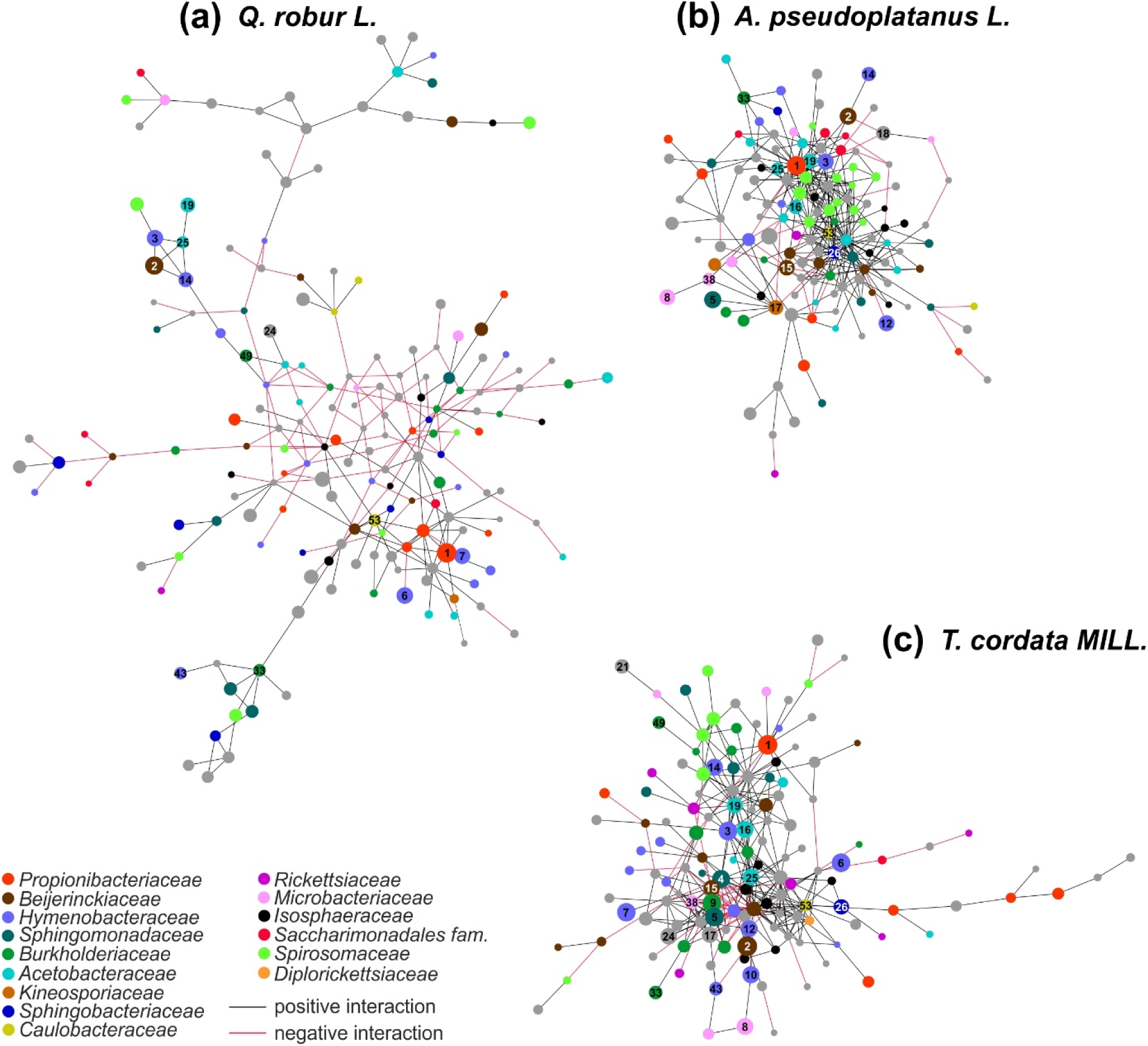
Co-occurrence network of bacterial species-level OTUs across height levels and individuals for each tree species. (a) *Q. robur L.*; (b) *A. pseudoplatanus L.*; (c) *T. cordata MILL*. Only network modules with more than 10 OTUs are shown. Color of circles denotes taxonomic affiliation on the family level. Numbers indicate core OTUs.

## Discussion

Phyllosphere microbiota in tree canopies play central roles in biogeochemical cycling and contribute to host plant fitness, protection and productivity (Laforest-Lapointe et al. 2019). Here, we hypothesized that both position in the canopy and tree species identity shape the phyllosphere microbial community in a floodplain hardwood forest in central Germany. In fact, we found evidence of an influence of both factors, however, canopy-position dependent effects were more pronounced and pointed to vertical gradients across the canopy. Bacterial abundance and OTU richness were lower at the top of the canopy compared to the mid canopy and bottom canopy positions across all three tree species – *Q. robur L., A. pseudoplatanus L., T. cordata MILL.*. While previous studies reported a large variation of phyllosphere bacterial community structure within a single tree canopy (Laforest-Lapointe et al. 2016a, Leff et al. 2015), trends of increasing diversity from the top towards the mid and bottom canopy have rarely been described for bacterial (Stone and Jackson 2019) or fungal communities (Harrison et al. 2016, Izuno et al. 2016, Christian et al. 2017). Here, we demonstrate that not only microbial diversity, but also microbial abundances follow the same trends and are likely affected by the same canopy position dependent factors.

Leaf-associated bacteria might simply be washed off with rainwater, leading to continuous loss of biomass and OTUs to the mid and bottom position of the canopy or eventually resulting in their transport to the soil via throughfall or stemflow (Bittar et al. 2018). In fact, Stone and Jackson (2019) found that rainfall influenced compositional similarity of bacterial communities throughout the canopy of *Magnolia* trees, which could also be one the mechanisms underlying the higher similarity between mid canopy and bottom canopy communities versus communities at the top of the canopy observed in our study. Similarly, interior canopy and upper canopy communities were the most distinct in the *Magnolia* canopy (Stone and Jackson 2019). However, these authors also reported that rain did not have any effect on bacterial species richness, questioning to which extent rainfall imposes physical disturbance on the phyllosphere-associated microbiota. Moreover, leaf-associated bacteria are protected against such physical forces by aggregates and biofilms and also by the surface structure of the leaves (Huber et al. 1997). Consequently, rainfall is likely not the main driver of lower abundance and diversity at the top of the canopy in our study.

Alternatively, more extreme environmental conditions, such as higher exposure to UV radiation, weather extremes, desiccation, and depletion of nutrients due to rain-mediated wash off, form a harsher environment for microbial colonization than the interior parts of the canopy (Stone and Jackson 2019). These conditions could lead to a selective enrichment of specialists at the top of the canopy, while the dominating phyllosphere bacteria of the mid and bottom canopy position become less competitive. Interestingly, Actinobacteria, in particular one OTU affiliated with the genus *Friedmaniella*, appeared to be the most responsive bacterial group to canopy position and showed a strong increase in its relative abundance from the top towards the mid and bottom canopy position. In general, distribution patterns of the 20 most abundant OTUs appeared to be strongly linked to canopy position, suggesting contrasting ecological preferences for bacteria related to *Friedmaniella* versus those related to *Hymenobacter, Methylobacterium, Kineococcus*, or *Massilia*, whose relative abundance increased at the top of the canopy. Previous findings suggested that general stress response is an essential mechanism for plant colonization by *Methylobacterium*, including responses to heat shock and desiccation, and oxidative, UV, ethanol and osmotic stresses (Gourion et al. 2006, 2008). In addition, increased abundances of *Methylobacterium* in upper parts of the canopy of *Magnolia* trees have been explained by a positive response of this genus to changes in leaf physiology following higher light or higher temperature, or by a direct response to these environmental parameters (Stone and Jackson 2019). In addition to a better adaption to harsh environmental conditions at the top of the canopy, increased relative abundances of *Methylobacterium, Hymenobacter, Kineococcus*, and *Massilia* may also have been supported by reduced abundances of *Friedmaniella* as a potentially very competitive inhabitant of the hardwood forest phyllosphere.

Plant species identity was identified as another key factor that influenced phyllosphere bacterial community composition in the floodplain forest, similar to tropical forests and temperate forest ecosystems in North America (Redford et al. 2010, Kembel et al. 2014, Laforest-Lapointe et al. 2017). Bacterial communities associated with oak and maple were more similar to each other than to those associated with lime trees, suggesting that oak and maple provided more favorable and more similar conditions for certain taxa, e. g., for Actinobacteria related to *Friedmaniella*, than did lime. Plant host attributes such as plant taxonomic identity and phylogeny, wood density, leaf mass per area, seed mass, leaf water content, and leaf nitrogen and phosphorus concentrations have been suggested as key factors underlying the relationship between plant species and their microbiome in neotropical as well as in temperate forests (Yadav et al. 2005, Kembel et al. 2014, Laforest-Lapointe et al 2016b). Additional factors may include the rate of production of volatile organic compounds such as methanol, which can act as important substrate for the phyllosphere microbiota (Westoby et al. 2002, Redford et al. 2010, Kim et al. 2012, Bringel and Coue 2015).

Co-occurrence network analysis revealed that the three tree species did not only differ in their bacterial community structure and OTU composition but also in the patterns how these OTUs were connected to each other across tree individuals, canopy position, and spatial replicates. The observed larger fraction of negative interactions between OTUs in association with oak may point to stronger vertical gradients within the oak canopy or larger variation across individuals of the same tree species. Moreover, OTUs with a central position in the network, e. g., by multiple connections to other OTUs, differed between tree species. These findings suggest that plant host-related factors and the chemical environment that they shape select for specific microbial core consortia that are strongly tree species dependent.

Interestingly, at a relative abundance of up to 50%, Actinobacteria were by far more prominent in our study compared to neotropical or tropical forests (Kembel et al. 2014, Kim et al. 2012) but also to previous investigations of Canadian temperate forests or the phyllosphere of hornbeam (*Carpinus betulus*), where they only accounted for 5 - 9% of the total community (Laforest-Lapointe et al. 2016b, Imperato et al. 2019). The most abundant OTU in our study was closely related to *Friedmaniella okinawensis* and *F. sagamiharensis* originally isolated from spider webs in a Japanese forest (Iwai et al. 2010). Bacteria related to *Friedmaniella* have been found in lower abundances in the phyllosphere of apple orchards or in urban environments (Yashiro et al. 2011, Espenshade et al. 2019) and can also grow as endophytes (Tuo et al. 2016, Pirttilä 2018). In fact, species within the genus *Friedmaniella* isolated from forest spider webs or the bark of mangrove plants have the capability to utilize a broader range of organic carbon compounds than other species of that genus (Iwai et al. 2010, Tuo et al. 2016), suggesting that this broader substrate spectrum could be one the mechanisms underlying their success in the phyllosphere.

Most of the other genus-level taxa representing the hardwood forest core microbiome, such as *Hymenobacter, Methylobacterium, Sphingomonas*, and *Pseudomonas* have frequently been observed in association with other temperate forest tree species (Laforest-Lapointe et al. 2016a, Stone and Jackson 2019) but also with herbaceous plants (Delmotte et al. 2009). The genus *Methylobacterium* uses methanol as its carbon and energy source, a C1 compound typically released by plants (Sy et al. 2005). Besides methanol, small amounts of nutrients, such as glucose, fructose, and sucrose (Lindow and Brandl 2003), but also amino acids, methane, terpenes, and chloromethane (Delmotte et al. 2009, Nadalig et al. 2011, Iguchi et al. 2012, Imperato et al. 2019) can leach from the interior of the plant and be available for the phyllosphere microbiota. Overall, our findings suggest that microbial processes in the hardwood forest canopies are largely dominated by heterotrophic or C1-dependent metabolisms. Although nitrification as previously been proposed as an important process in tree canopies, stimulated by excess atmospheric deposition of ammonia (Papen et al. 2002, Guerrieri et al. 2015), we found only few sequence reads affiliated with chemolithoautotrophic *Nitrosomonadaceae*.

Given the temporal development of forest tree canopies throughout the growing season, our sampling provides only one snapshot, and the extent to which the September phyllosphere communities differ from earlier stages in spring and summer remains currently unclear. A previous study in a temperate mixed forest showed that temporal effects were smaller than those associated with host species identity (Laforest-Lapointe et al. 2016b). However, we cannot rule out that the high relative abundance of *Friedmaniella* could be linked to a stage of early senescence of the leaves. Successional changes in phyllosphere communities can be associated with changes in the physiology of the host plant but can also be shaped by the constant import of microbes from various sources such as air, soils, rainwater, and animal and plant dispersal vectors (Kembel et al. 2014, Bai et al. 2015, Sanchez-Canizares et al. 2017). Representatives of the genera *Hymenobacter, Methylobacterium, and Massilia* have been reported from air samples (Zhen et al. 2018) or from aerosols originating from agricultural practices (Rastogi et al. 2012, Bringel and Coue 2015), suggesting that airborne microbes play a major role in the early colonization of the surfaces of young leaves in spring (Bringel and Coue 2015) and could continuously be introduced to the phyllosphere communities throughout the season. Consequently, the September phyllosphere represents a stage that integrates the results of different mechanisms of colonization and competitive interactions between phyllosphere microbiota throughout the growing season.

## Conclusions

Our findings clearly demonstrate that both position in the canopy and tree species have a strong effect on the structure of phyllosphere bacterial communities in a floodplain hardwood forest. Consistently lower bacterial diversity at the top of the canopy compared to the canopy mid and bottom positions pointed to a stronger selective pressure on phyllosphere bacteria given presumably harsher environmental conditions at the treetop. Across all three tree species, we observed a striking predominance of Actinobacteria related to *Friedmaniella* sp., which could be a typical feature of floodplain hardwood forests or linked to the early senescent state of leaves sampled in mid September.

## Acknowledgements

We thank Rolf Engelmann for technical support during sampling and Christian Wirth for providing access to the Leipzig canopy crane facility. Julia Rosenberger is acknowledged for help with sample processing. Sequencing was financially supported by the German Center for Integrative Biodiversity Research (iDiv) – Halle, Jena, Leipzig funded by the Deutsche Forschungsgemeinschaft (FZT 118). Additional support was provided by the Collaborative Research Centre AquaDiva (CRC 1076 AquaDiva) of the Friedrich Schiller University Jena, funded by the Deutsche Forschungsgemeinschaft.

