## Supplemental_Information for "Canopy position has a stronger effect than tree species identity on phyllosphere bacterial diversity in a floodplain hardwood forest"

Friedrich Schiller University Jena

Institute of Biodiversity – Aquatic Geomicrobiology

Dornburger Strasse 159

D-07743 Jena

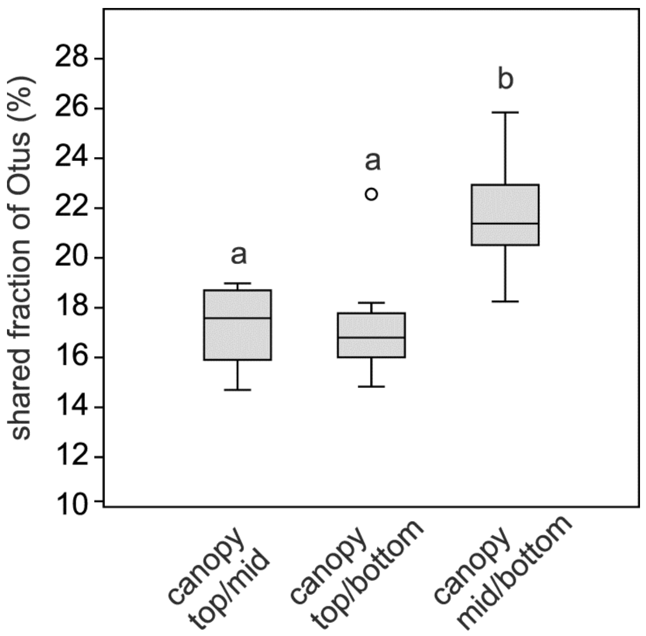


**Supplementary Fig. 1:**

Fraction of OTUs shared between the canopy's top and mid position, top and bottom position, and mid and bottom position, respectively across all three tree species. Box plots represent mean values calculated from three spatial replicates per tree individual and height within the canopy.

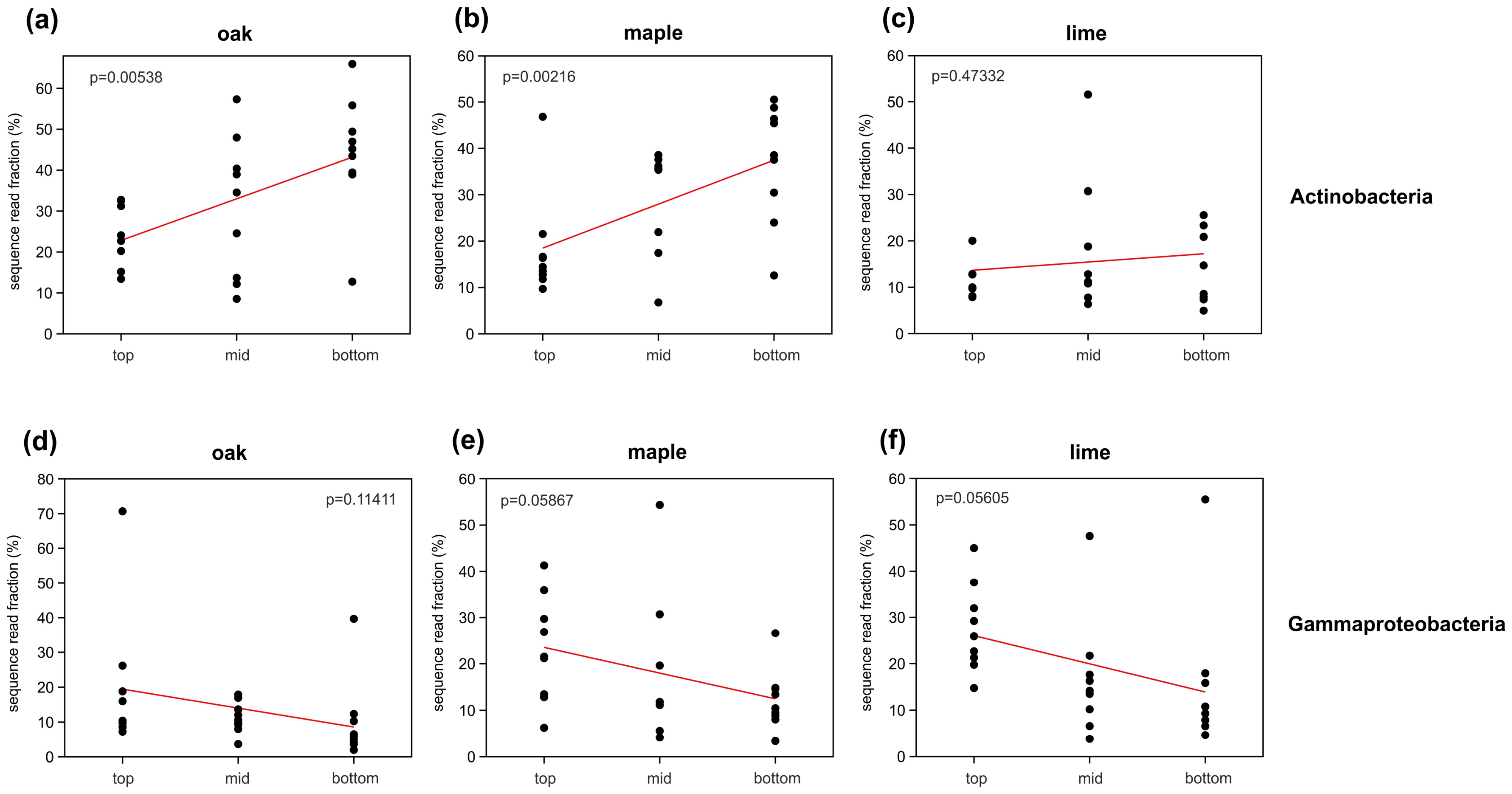


**Supplementary Fig. 2**

Linear regression of changes in relative abundances of Actinobacteria (a-c) and Gammaproteobacteria (d-f) from the bottom position of the canopy via mid position to the top of the canopy. Analyses were performed separately for each tree species. Data points originate from three individuals of a given tree species with three spatial replicates per position in the canopy.

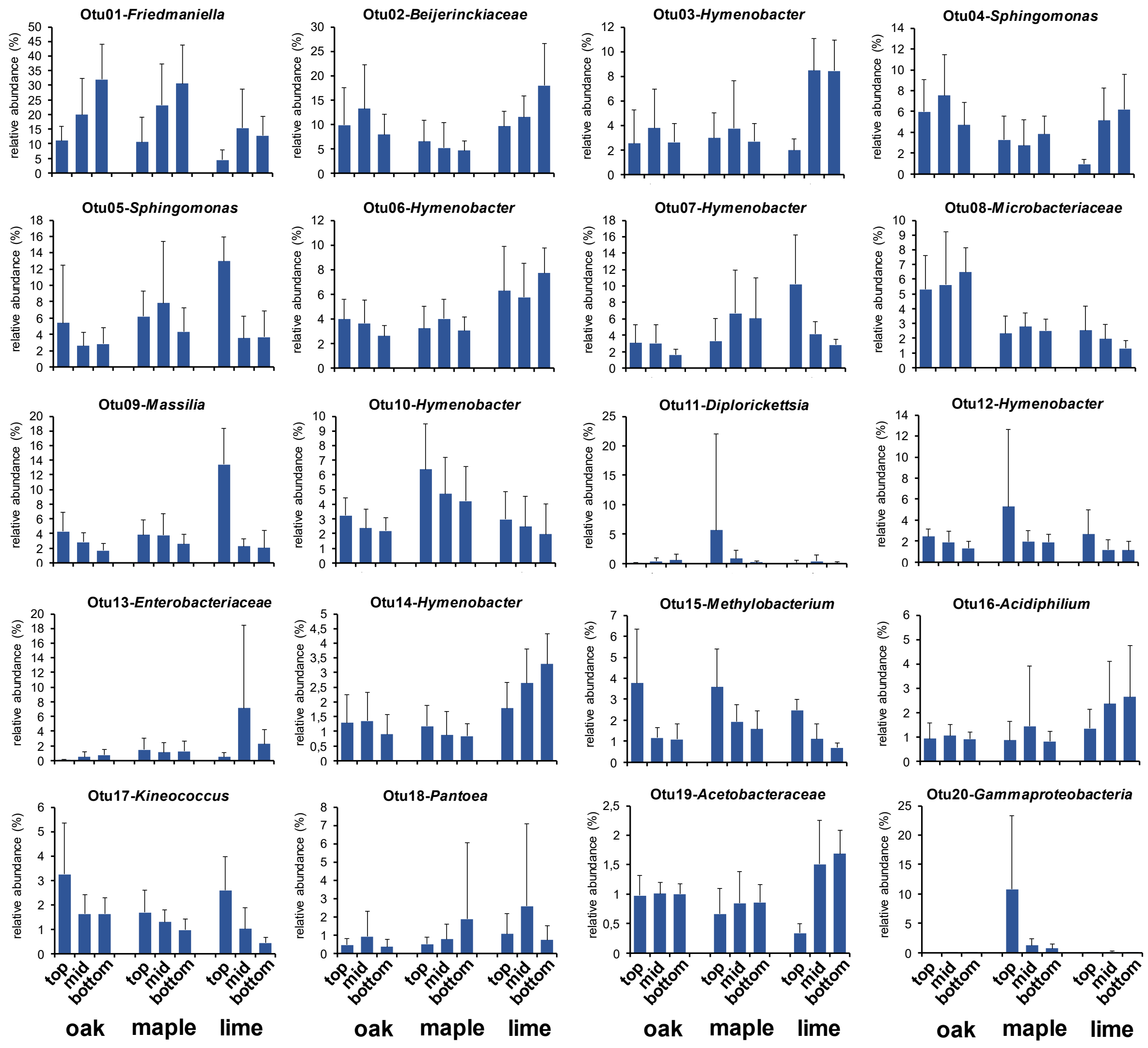


**Supplementary Fig. 3:**

Changes in relative abundances of the first 20 most abundant OTUs across bottom, mid, and top canopy positions, shown separately for the three tree species. Data are mean (± standard deviation) of three individuals of a tree species, sampled in three spatial replicates per canopy position.

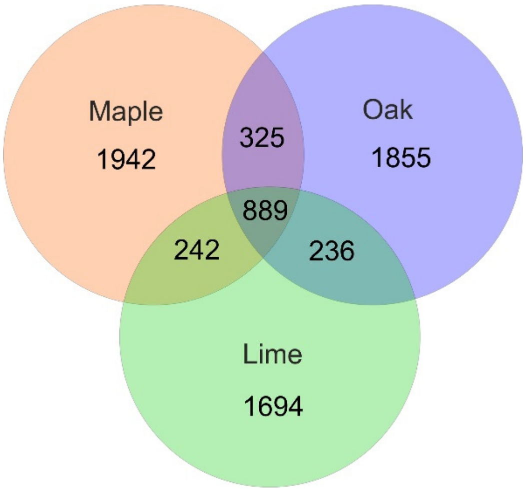


**Supplementary Fig. 4:**

Number of species-level OTUs shared between tree species. OTUs associated with maple, oak, and lime across individuals, canopy positions, and spatial replicates were merged to one OTU pool per tree species, and comparisons of shared OTUs were made between these tree-species specific OTU pools.

**Supplementary Table 1:**

Abundances of bacterial 16S rRNA genes per g (dry weight) in association with leaves of oak, maple, and lime at top, mid, and bottom canopy positions. Data are means and standard deviation (in parentheses) of three technical qPCR replicates. For a number of samples, leaf dry weight is not available due to loss of plant material after DNA extraction and thus, gene abundances could not be determined (n. a.).

| tree species | canopy position | sample | | | | | | | | |
|---|---|---|---|---|---|---|---|---|---|---|
| | | **Qr7** | **Qr8** | **Qr9** | **Qr16** | **Qr17** | **Qr18** | **Qr25** | **Qr26** | **Qr27** |
| oak | top | n. a. | n. a. | 3.1 x $10^{7}$ (2.2 x $10^{6}$) | n. a. | n. a. | 6.18 x $10^{6}$ (9.6 x $10^{4}$) | n. a. | n. a. | 1.4 x $10^{7}$ (3.6 x $10^{6}$) |
| | | **Qr4** | **Qr5** | **Qr6** | **Qr13** | **Qr14** | **Qr15** | **Qr22** | **Qr23** | **Qr24** |
| oak | mid | n. a. | 7.0 x $10^{7}$ (2.0 x $10^{6}$) | 6.4 x $10^{7}$ (2.0 x $10^{6}$) | 1.9 x $10^{8}$ (2.2 x $10^{6}$) | 1.5 x $10^{8}$ (6.8 x $10^{6}$) | 1.4 x $10^{8}$ (3.1 x $10^{6}$) | 6.2 x $10^{7}$ (1.6 x $10^{7}$) | 8.9 x $10^{7}$ (9.9 x $10^{6}$) | 2.8 x $10^{8}$ (2.5 x $10^{7}$) |
| | | **Qr1** | **Qr2** | **Qr3** | **Qr10** | **Qr11** | **Qr12** | **Qr19** | **Qr20** | **Qr21** |
| oak | bottom | 1.1 x $10^{8}$ (1.0 x $10^{7}$) | 2.2 x $10^{8}$ (1.4 x $10^{8}$) | 2.0 x $10^{8}$ (7.3 x $10^{7}$) | 9.9 x $10^{7}$ (6.5 x $10^{6}$) | 2.5 x $10^{8}$ (2.1 x $10^{7}$) | 2.4 x $10^{8}$ (2.3 x $10^{7}$) | n. a. | n. a. | n. a. |
| | | **Ap7** | **Ap8** | **Ap9** | **Ap16** | **Ap17** | **Ap18** | **Ap25** | **Ap26** | **Ap27** |
| maple | top | n. a. | n. a. | n. a. | n. a. | n. a. | 3.1 x $10^{8}$ (3.0 x $10^{7}$) | n. a. | n. a. | 9.8 x $10^{7}$ (7.6 x $10^{6}$) |
| | | **Ap4** | **Ap5** | **Ap6** | **Ap13** | **Ap14** | **Ap15** | **Ap22** | **Ap23** | **Ap24** |
| maple | mid | 4.2 x $10^{8}$ (6.7 x $10^{6}$) | 1.1 x $10^{8}$ (2.2 x $10^{6}$) | 1.1 x $10^{8}$ (6.3 x $10^{6}$) | 3.8 x $10^{8}$ (2.1 x $10^{7}$) | 9.6 x $10^{8}$ (6.1 x $10^{7}$) | 6.9 x $10^{8}$ (5.6 x $10^{7}$) | 9.5 x $10^{8}$ (2.6 x $10^{7}$) | 4.5 x $10^{8}$ (2.3 x $10^{7}$) | 6.2 x $10^{8}$ (4.5 x $10^{7}$) |
| | | **Ap1** | **Ap2** | **Ap3** | **Ap10** | **Ap11** | **Ap12** | **Ap19** | **Ap20** | **Ap21** |
| maple | bottom | 2.5 x $10^{8}$ (8.8 x $10^{6}$) | n. a. | n. a. | 2.9 x $10^{8}$ (4.3 x $10^{7}$) | n. a. | n. a. | 8.3 x $10^{8}$ (3.7 x $10^{7}$) | n. a. | 6.4 x $10^{8}$ (5.0 x $10^{7}$) |

**Supplementary Table 1, continued**

| tree species | canopy position | sample | | | | | | | | |
|---|---|---|---|---|---|---|---|---|---|---|
| | | **Tc7** | **Tc8** | **Tc9** | **Tc16** | **Tc17** | **Tc18** | **Tc25** | **Tc26** | **Tc27** |
| lime | top | n. a. | n. a. | 1.6 x $10^8$ (9.8 x $10^6$) | n. a. | n. a. | 5.2 x $10^8$ (4.0 x $10^7$) | n. a. | n. a. | 2.9 x $10^8$ (8.6 x $10^6$) |
| | | **Tc4** | **Tc5** | **Tc6** | **Tc13** | **Tc14** | **Tc15** | **Tc22** | **Tc23** | **Tc24** |
| lime | mid | 4.8 x $10^8$ (1.3 x $10^7$) | 3.6 x $10^8$ (2.1 x $10^7$) | 2.9 x $10^8$ (1.6 x $10^7$) | n. a. | 9.3 x $10^8$ (4.7 x $10^7$) | 7.0 x $10^8$ (2.6 x $10^7$) | 3.2 x $10^9$ (1.6 x $10^8$) | 2.1 x $10^9$ (1.6 x $10^8$) | n. a. |
| | | **Tc1** | **Tc2** | **Tc3** | **Tc10** | **Tc11** | **Tc12** | **Tc19** | **Tc20** | **Tc21** |
| lime | bottom | 4.8 x $10^8$ (4.3 x $10^7$) | 2.8 x $10^8$ (4.5 x $10^6$) | 2.2 x $10^8$ (6.3 x $10^6$) | 7.4 x $10^8$ (1.7 x $10^7$) | 1.1 x $10^9$ (5.4 x $10^7$) | n. a. | 3.2 x $10^9$ (1.6 x $10^8$) | 2.1 x $10^9$ (1.6 x $10^8$) | 5.2 x $10^8$ (4.0 x $10^7$) |
